# A location-specific spreadsheet for estimating Zika risk and timing for Zika vector surveillance, using U.S. military facilities as an example

**DOI:** 10.1101/088823

**Authors:** Desmond H. Foley, David B. Pecor

## Abstract

Local Zika virus transmission in the United States involving one or both of the known vector species, *Aedes aegypti* and *Ae. albopictus,* is of major concern. To assist efforts to anticipate the risks of transmission, we developed an Excel spreadsheet tool that uses vector and virus temperature thresholds, remotely sensed maximum temperature, and habitat suitability from models to answer the questions: “is Zika transmission likely here?” and “when should we conduct vector surveillance?”. An example spreadsheet, updated regularly and freely available, uses near real-time and forecast temperature data to generate guidance, based on a novel four level Zika risk code, for 733 U.S. military facilities in the 50 states, the District of Columbia, and the territories of Guam and Puerto Rico.

## INTRODUCTION

In 2016, Zika virus disease and congenital infections became nationally notifiable conditions in the United States (Council of State and Territorial Epidemiologists, 2016). A total of 2,382 confirmed and probable cases of ZIKAV disease with illness onset were reported to ArboNET, the U.S. national arboviral surveillance system managed by CDC and state health departments, during January 1 – July 31, 2016 (Walker et al. 2016). In July 2016 the first locally acquired cases of Zika virus (ZIKAV) from mosquitoes were confirmed for the U.S. state of Florida (Likos, 2016). *Aedes* mosquitoes transmit ZIKAV, chikungunya virus (CHIKV), dengue virus (DENV), and yellow fever virus (YFV), among others, so co-infections are possible. Although *Ae. albopictus* is thought to be a competent vector of ZIKAV (Grard et al. 2014), *Ae. aegypti* has been implicated as the primary transmitter of the virus in human populations in the ongoing outbreak in the Americas (Guerbois et al. 2016, Ferreira-de-Brito et al. 2016). This is likely the result of *Ae. aegypti* preferring to feed more frequently on humans (Scott et al. 1993, 2000), and being highly peridomestic compared to *Ae. albopictus*, which can inhabit more rural environments (Braks et al. 2003; Tsuda et al. 2006).

In this study, we concentrated on U.S. Department of Defense (DoD) facilities but the approach could be used for any area of interest. Some military facilities have long standing mosquito surveillance programs (Foley et al. 2011a), and Zika virus surveillance is being enhanced in the U.S. military as a result of the recent threat, for example, through funding from the Global Emerging Infections Surveillance and Response (GEIS), section of the Armed Forces Health Surveillance Branch in the Defense Health Agency’s Public Health Division (Pellerin, 2016). According to a March 2016 U.S. DoD memo, 190 DoD installations are located in areas where mosquitoes capable of carrying ZIKAV occur, and increased vector monitoring will be conducted in installations in 27 states, the District of Columbia, Guam and Puerto Rico (Kime, 2016). Four regional commands exist under the U.S. Army Medical Command, and all of these have Entomological Sciences Divisions that conduct mosquito surveillance. Additionally, the U.S. Air Force School of Aerospace Medicine, the U.S. Navy and Marine Corps Public Health Center and regional Navy Environmental and Preventative Medicine Units, and the Navy Entomology Center of Excellence, assist those undertaking vector surveillance or arbovirus testing.

For a military entomologist tasked with establishing and maintaining an *Aedes* spp. / ZIKAV surveillance program in temperate areas that experience high mosquito seasonality, two important questions arise: 1) is ZIKAV transmission possible here?; and 2) when should we conduct vector surveillance?. In the following we describe an Excel-based tool that is designed to assist entomologists and other health personnel address these two questions.

Habitat suitability models displaying potential distribution have been published for both *Ae. aegypti* and *Ae. albopictus* (Attaway et al. 2016, Brady et al. 2014, Campbell et al. 2015, Khormi & Kumar 2014, Medley 2010), as well as for ZIKAV (Carlson et al. 2016, Messina et al. 2016, Perkins et al. 2016, Samy et al. 2016). While these models often display average yearly suitability they do not necessarily provide information that could be used for decisions about the timing of surveillance activities, and are global in extent rather than focused on particular areas where a surveillance program might be established. Questions about timing of mosquito monitoring and allocation of resources requires a consideration of what conditions limit adult mosquito activity and ZIKAV dissemination in the field.

Relative humidity, rainfall, drought, and wind velocity affect survival and behavior of mosquitoes, and therefore transmission (Kramer & Ebel, 2003). However, temperature is the most important ecological determinant of development rate in *Ae. aegypti* (Couret & Benedict 2014), and one of the principal determinants of *Aedes* survival (Brady et al. 2013). Temperature also directly affects the replication rate of arboviruses, thus affecting the extrinsic incubation period (Gubler et al. 2007). What then, do we know about how temperature limits *Aedes* and arboviruses like ZIKAV?

In Saudi Arabia, Khormi et al. (2011) found that the minimum temperature range of 18-25 °C is suitable for *Ae. aegypti* survival, and the survival rate increases up to 38 °C. Conner (1924) and Wayne & Graham (1968) found that *Ae. aegypti* is most active at temperatures between 15 °C and 30 °C, while other field and laboratory observations found survival rates from about 18 °C to ≤ 38 °C, based on daily or monthly minimum and maximum temperatures (Macfie, 1920; Bliss & Gill, 1933; Christopher, 1960). In a study of *Ae. aegypti* distribution using the program CLIMEX, Khormi & Kumar (2014) set the limiting low temperature at 18 °C, the lower optimal temperature at 25 °C, the upper optimal temperature at 32 °C and the limiting high temperature at 38 °C. Brady et al. (2014) limited their predictions of temperature suitability to areas with a maximum monthly temperature exceeding 13°C for *Ae. albopictus* and 14°C for *Ae. aegypti*. These threshold temperatures were based on previous studies of the observed temperatures below which biting and movement behaviors are impaired [Christophers, 1960; Estrada-Franco & Craig, 1995; Carrington et al. 2013a,b).

Studies suggest that an increase between 14-18 °C and 35-40 °C can lead to higher transmission of dengue (Wallis, 2005). Xiao et al. (2014) found that oral infections of DENV2 did not produce antigens in the salivary glands of *Ae. albopictus* kept at 18°C for up to 25 days but did produce antigens at 21°C during this period. It is not known if *Ae. albopictus* held longer at the lower temperature would have disseminated infections, but Dohm et al. (2002) found that *Culex pipiens* required 25 days at 18°C to disseminate infections of West Nile Virus. For comparison, WNV is capable of replication from 14-45°C (Cornel et al. 1993, Kinney et al. 2006). Tilston et al. (2009) analyzed monthly average temperature of cities that experience chikungunya outbreaks and found that start and finish occurred when average monthly temperatures were 20°C or higher. At the upper temperature limit, Kostyuchenko et al. (2016) found that ZIKAV is more thermally stable than DENV, and is also structurally stable even when incubated at 40°C, mimicking the body temperature of extremely feverish patients after virus infection (but see Goo et al. 2016).

Remotely sensed temperature data is freely available from multiple sources as both near-real time recordings and forecast predictions. Combining remotely sensed temperature data with predicted distributions of the vectors and virus could provide insight into when areas of interest are suitable for transmission and should be actively monitored. Our aim was to produce a knowledge product and surveillance decision tool that makes use of publicly available information about potential distribution and thermal requirements of the vectors and virus at U.S. military facilities.

## MATERIALS AND METHODS

### Areas of interest

The location and boundary of U.S. military facilities was obtained from the US Census Bureau’s TIGER/Line 2015 shapefile product (<u>http://www.census.gov/geo/maps-data/data/tiger.html</u>). This shapefile lists facilities in the continental United States (CONUS), Alaska, Hawaii, Puerto Rico and Guam. As some facility names comprised multi-part polygons, these were reduced from 804 to 733, to match the number of unique facility names, using the Dissolve tool in ArcMap 10.4 (ESRI, Redmond, CA – used throughout). The centroid of each facility was selected to produce a shapefile of points using the Feature to Point tool (inside polygon option checked) of ArcMap. The georeference of each point was obtained by the Add XY Coordinates tool and joined to the points shapefile. Extraction of all facility centroid raster values was first obtained by the Extract values to points tool then for polygons using the Zonal statistics as Table tool, and the results merged. This approach was needed because smaller polygons would not produce results using the Zonal statistics as Table tool, which necessitated using the raster data associated with the points for these facilities.

### Temperature data

To monitor temperature in near real-time, daily time averaged maps of air temperature at the surface (Daytime/Ascending) were downloaded from the Giovanni 4.19 (Released Date: 2016-04-12. Data provided by the NASA Goddard Earth Sciences (GES) Data and Information Services Center (DISC)) data portal at 1° spatial resolution. Daily gridded temperature analyses were also collected from the NOAA, U.S. National Weather Service Climate Prediction Center (CPC). Forecast temperature data was also provided by the CPC and the NOAA National Digital Forecast Database (NDFD) at 5 km spatial resolution. For predictions based on monthly averages, monthly gridded climate data with a spatial resolution of 1 km were downloaded from the WorldClim (Version 1.4) Global Climate Data center.

### Habitat suitability models

We chose the models of *Ae. aegypti* and *Ae. albopictus* by Kraemer et al. (2015), as these are recent and are based on an extensively documented set of presence observations for each vector. For this study, we used the habitat suitability model for ZIKAV transmission by Messina et al. (2016). The 0.5 model suitability score was arbitrarily used as the cut-off for presence/absence.

### Thresholds

Temperatures suitable for activity of *Ae. aegypti* and *Ae. albopictus* combined was estimated as 13 – 38°C, and for ZIKAV this was 18 – 42°C. We acted conservatively by using temperatures at the extremes of the reported suitable temperature range, and maximum rather than mean air temperatures.

### Human population data

In order to more fully understand the potential impact of ZIKAV risk to military and non-military personnel and their families in and around each facility, we explored risk in terms of human population data, with the following considerations. The flight range of *Ae. aegypti* and *Ae. albopictus* is in the order of hundreds of meters only (Honório et al. 2003, Harrington et al. 2005) and each facility would differ in the average distance that human carriers of ZIKAV would routinely travel to and from each facility. Additionally, some facilities are remote, while others are adjacent to or enclosed within urban and suburban areas. Despite these complications, we created a buffer of 5 km around all facility polygons to capture the human population density according to LandScan 2011 (Oak Ridge National Laboratory). This was accomplished using the LandScan raster and the Buffer and sum output in the Zonal Statistics as Table tools in ArcMap. A buffer of 5 km is a conservative estimate and is meant to give a uniform measure for each facility of the potential host density effected in an outbreak or vector control situation.

### Excel-based Zika risk tool

A goal of this project was to display disparate data sources visually and in a simple and intuitive way in order to more effectively communicate the level of risk at each military facility. The risk estimation and alert system needed to be in a format that was readily understandable and that can be easily accessed by military users, who often have IT security restrictions or bandwidth caps. We chose MS-Excel^®^ (Microsoft Corp, Seattle, WA), as a universal platform for performing calculations and reporting results. This software had the added advantage that the scatterplot function can be used to map each military facility (Foley, 2011b), with icons displaying various categories of risk, and using a geocorrected map background (Esri, DeLorme, USGS, NPS. World Terrain Base - Sources: Esri, USGS, NOAA) for each U.S. State. Other notable features that were used in the Excel risk estimation tool were the formula functions, conditional formatting to represent categories of numbers as different types of symbols, dependent dropdown lists and hyperlinks to allow users to navigate more quickly to the results of individual facilities, and textualized results that users can read as statements describing the situation and as guidance for vector surveillance.

### Calculations within the Excel Risk Estimation Tool

Given the maximum temperature is available for a site (i.e. “Temp.”), the following lists an example sequence of tasks and their calculations, with explanations and the Excel formula (in square brackets), culminating in a risk rating:

1. Column A. “Was temperature suitable during period for the vector?”, i.e. if the maximum was 13 to 38°C, it is 1 otherwise 0 [ =IF((Temp.>=13)-(Temp.>38),1,0)],
2. Column B. “Was temperature suitable during period for virus replication in mosquito?”, i.e. if the maximum was 18 to 42°C, it is 1 otherwise 0 [=IF((Temp.>=18)-(Temp.>42),1,0)]
3. Column C. What is the sum of the thermal suitability values for vector (Column A) and virus (Column B)? (i.e. possible choices are: 0, 1 or 2)
4. Column D. If temperature for the vectors (Column A) was within the required range, what is the model suitability for vector?, i.e. this was the maximum modeled suitability (0 – 1.00) for either *Ae. aegypti* or *Ae. albopictus*
5. Column E. Score vector model suitability as 3 if >=0.5, otherwise 2 [=IF(Column D<0.5,2,IF(Column D>=0.5,3))]. The 0.5 model suitability score was arbitrarily used as the cut-off for presence/absence.
6. Column F. If temperature for the virus (Column B) was within the required range, what is the model suitability for the virus? (0 – 1.00)
7. Column G. Score virus model suitability as 7 if >=0.5, otherwise 5 [=IF(Column F<0.5,5,IF(Column F>=0.5,7))]. The 0.5 model suitability score was arbitrarily used as the cut-off for presence/absence.
8. Column H. What is the “Combined Score” for the interaction of temperature suitability of vector and virus, vector model suitability, and virus model suitability score (i.e. = C*E*G)? The use of 0, 1 and prime numbers for the component scores produces a unique semi-prime number for the product, i.e. 0, 10 (=1*2*5), 14 (=1*2*7), 15 (=1*3*5), 21 (=1*3*7), 20 (=2*2*5), 28 (=2*2*7), 30 (=2*3*5), or 42 (=2*3*7). A zero indicated that the temperature at the site was unsuitable for both vector and pathogen, so suitability scores were irrelevant, and combination scores were all scored zero.
9. The nine possible Combined Scores were divided into 6 categories based on whether preconditions do not exist for transmission, are unsuitable for transmission, are somewhat suitable for transmission, or are suitable for transmission (Figure 1).
10. Column I. An Overall Zika Risk Code was established based on the Combined Score (Figure 1), and rates conditions as low (Blue: Code 1) to high risk (Red: Code 4). The nine possible Combined Scores are initially divided according to whether temperature conditions are not suitable (Code 1), or suitable, for the vectors and virus (Codes 2 - 4). Codes 2 - 4 are then characterized according to increasing habitat suitability, with Code 4 being where models predict suitable habitat for vectors and virus.
11. Action statements were constructed based on the temperature and habitat model suitability scores (Figure 2). For example, if conditions are too cold for the development of the vectors, and models predict that the location is unsuitable for the vectors then the Action statement would be: “Too cold or hot for vectors - surveillance unnecessary. When temp. suitable, model suggests vectors unlikely or low numbers.”. Alternatively, if conditions are warm enough for the development of the vectors, and models predict that the location is highly suitable for the vectors then the Action statement would be: “Temp. suitable for vectors - surveillance may be needed. When temp. suitable, model suggests vectors likely - may need control, education, and policies minimizing exposure”.

**Figure 1.**
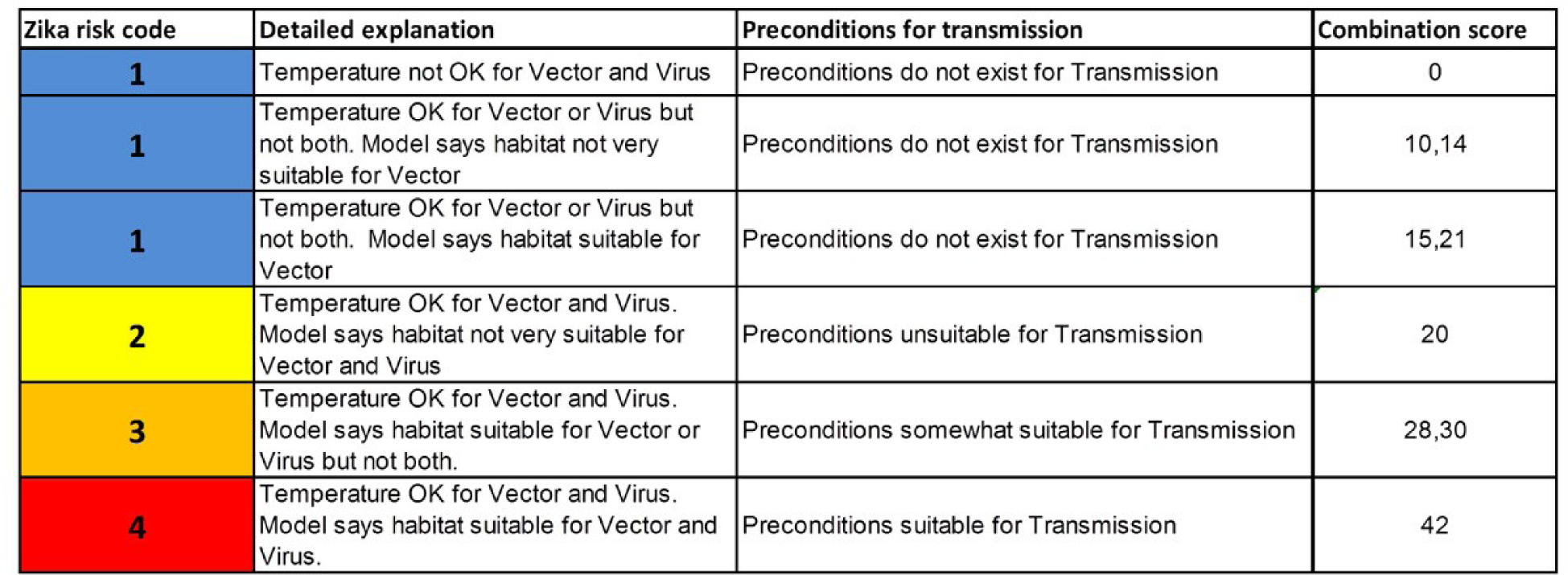
An overall Zika Risk Code based on the combined score, which rates conditions from low risk (Blue: Code 1) to high risk (Red: Code 4).

**Figure 2.**
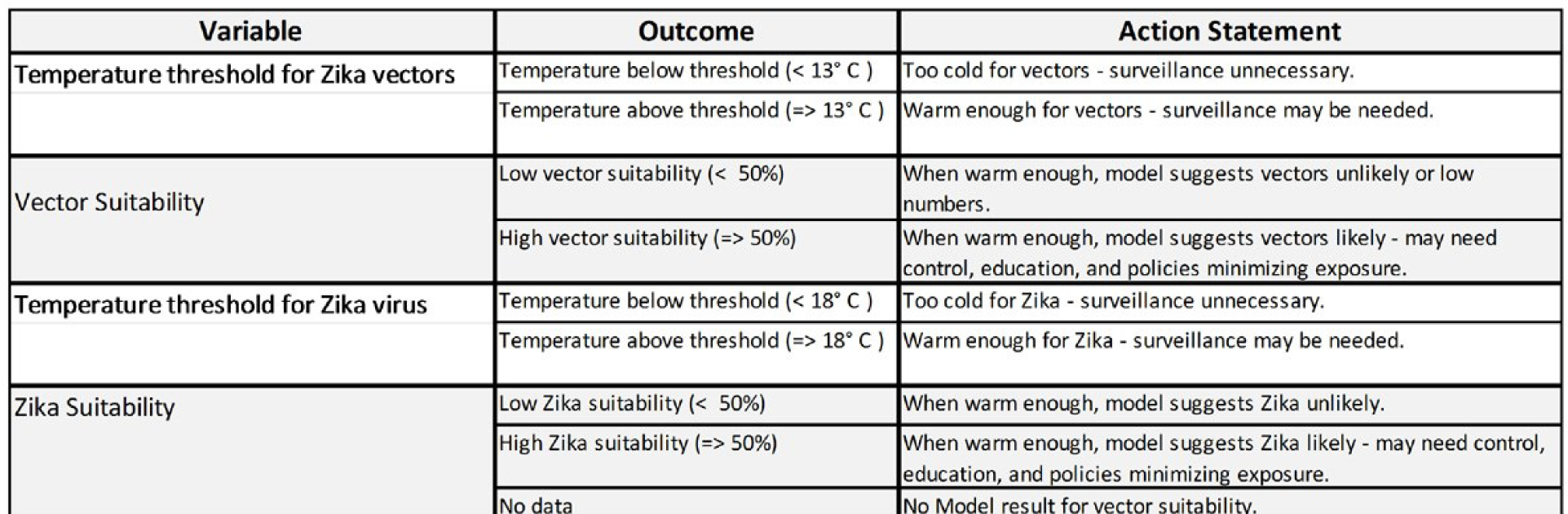
Action statements constructed on the basis of the temperature and habitat model suitability scores

## RESULTS

The Excel files provided comprise 12 monthly files based on average monthly maximum temperatures (suitable for longer term planning), and near real-time and forecast file, updated weekly. These files are freely available via the VectorMap website (http://vectormap.si.edu/Project_ESWG_ExcelZika.htm). The tool provides risk maps of facilities as a continental overview (Figure 3), and on a U.S. State basis (Figure 4). Results for individual facilities are navigable via dropdown menus and hyperlinks (Figure 5). State-wide summary data of risk profile and humans potentially impacted is given in Figure 6. The temporal changes in average risk based on the 12 monthly files is given in Figure 7 in terms of the number of facilities affected (of 733) and the number of people within 5 km of these facilities. April to October was the period of greatest risk with suitable conditions for Zika transmission (i.e. code 4) potentially affecting a maximum of 114 facilities in 12 states and territories, and 4,546,505 people within the vicinity of these facilities, of a total of 32,811,618 within the vicinity of all 733 facilities. The maximum number of facilities recording code 4 in any one month (e.g. August) were: Florida (36), Hawaii (16), Louisiana (12), Texas (11), and Virginia (11). Of these, the number of people within 5 km of these facilities were: Texas (1,215,230), Florida (1,125,032), Louisiana (462,586), Virginia (409,066), and Hawaii (247,918).

**Figure 3.**
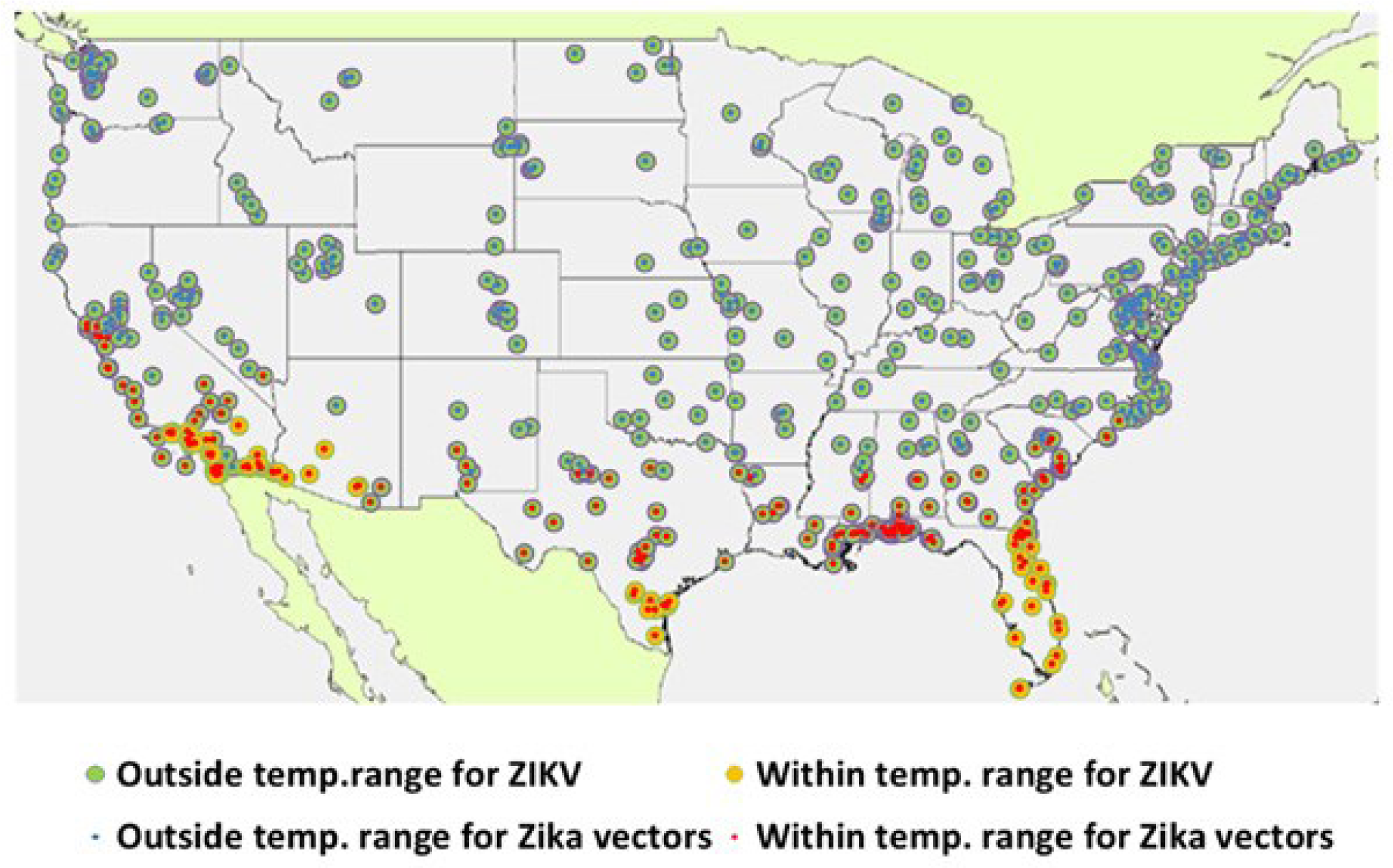
Average maximum temperature conditions for January for vectors and ZIKAV at military facilities within the lower 48 states of the U.S.

**Figure 4.**
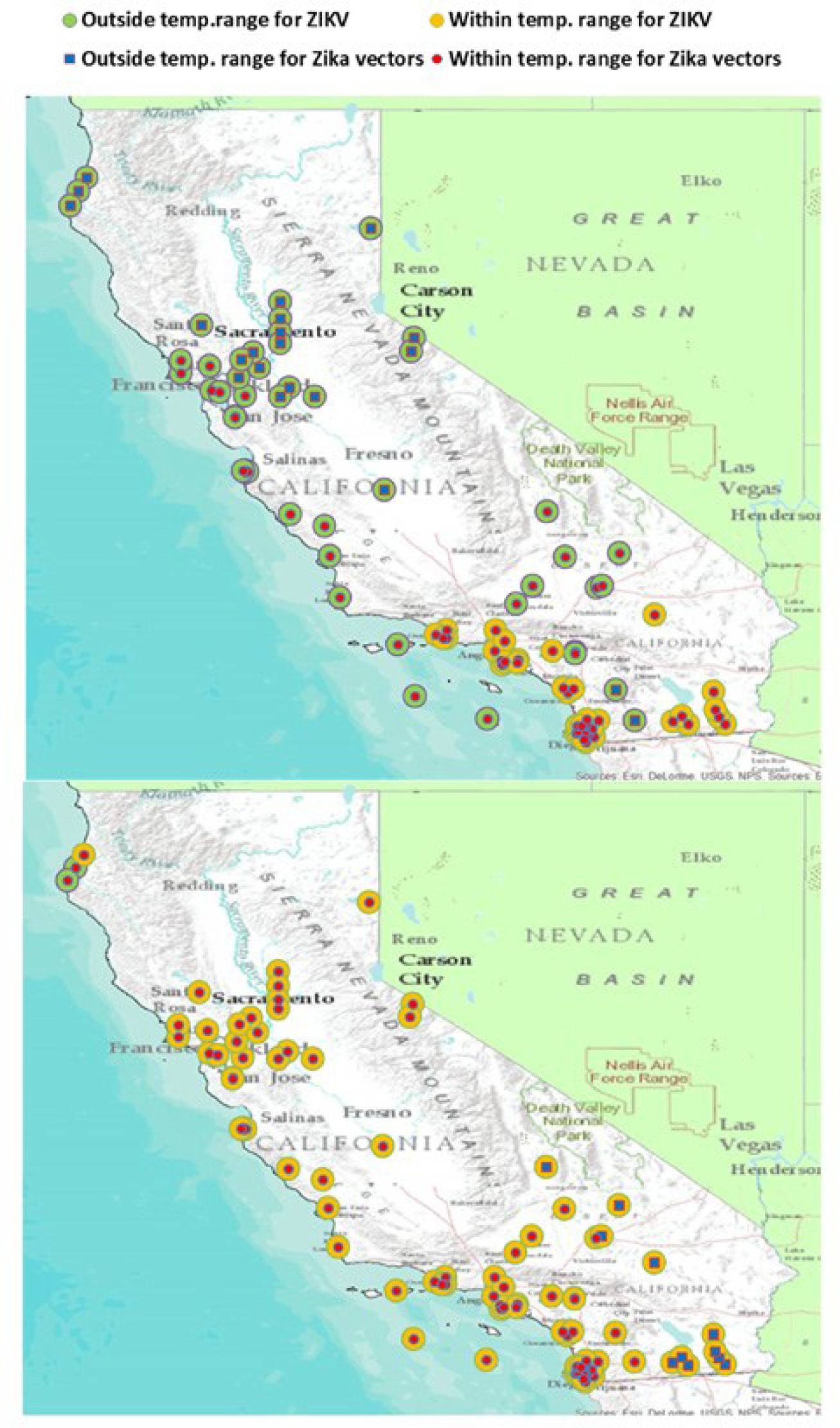
Thermal conditions for January (above) and August (below) for vectors and ZIKAV at military facilities in California. Note, unsuitable conditions in August in the south are due to temperatures being too high for the vectors.

**Figure 5.**
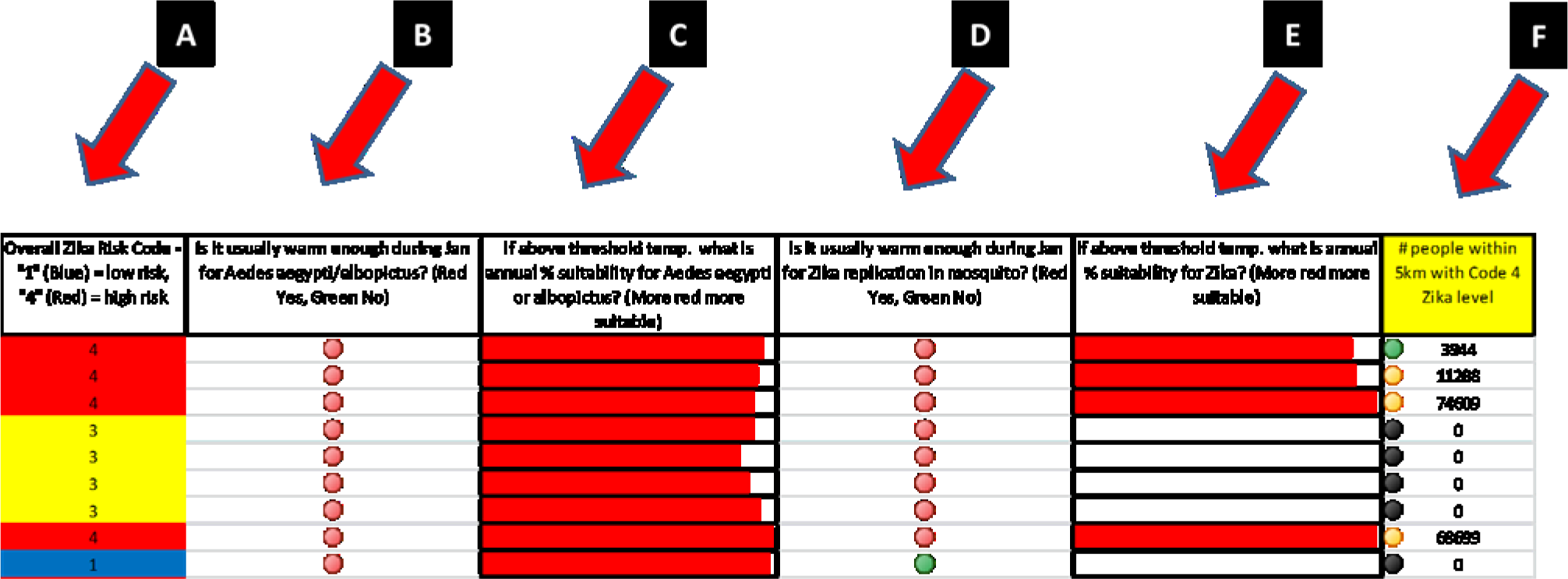
An overall Zika virus risk code for near real-time and forecast periods (A) is assigned based on a combination of: temperature suitability for adult activity of the vectors (*Ae. aegypti* and *Ae. albopictus*) (B), modeled habitat suitability of the vectors (C), temperature suitability for Zika virus replication within the vectors (D), and modeled habitat suitability of Zika virus transmission (E). If suitable conditions exist (Zika Risk Code 4), the number of people within 5 km is shown (F) as one indication of the number of potential hosts in the vicinity, or the number of humans potentially benefiting from facility-wide vector surveillance and control programs.

**Figure 6.**
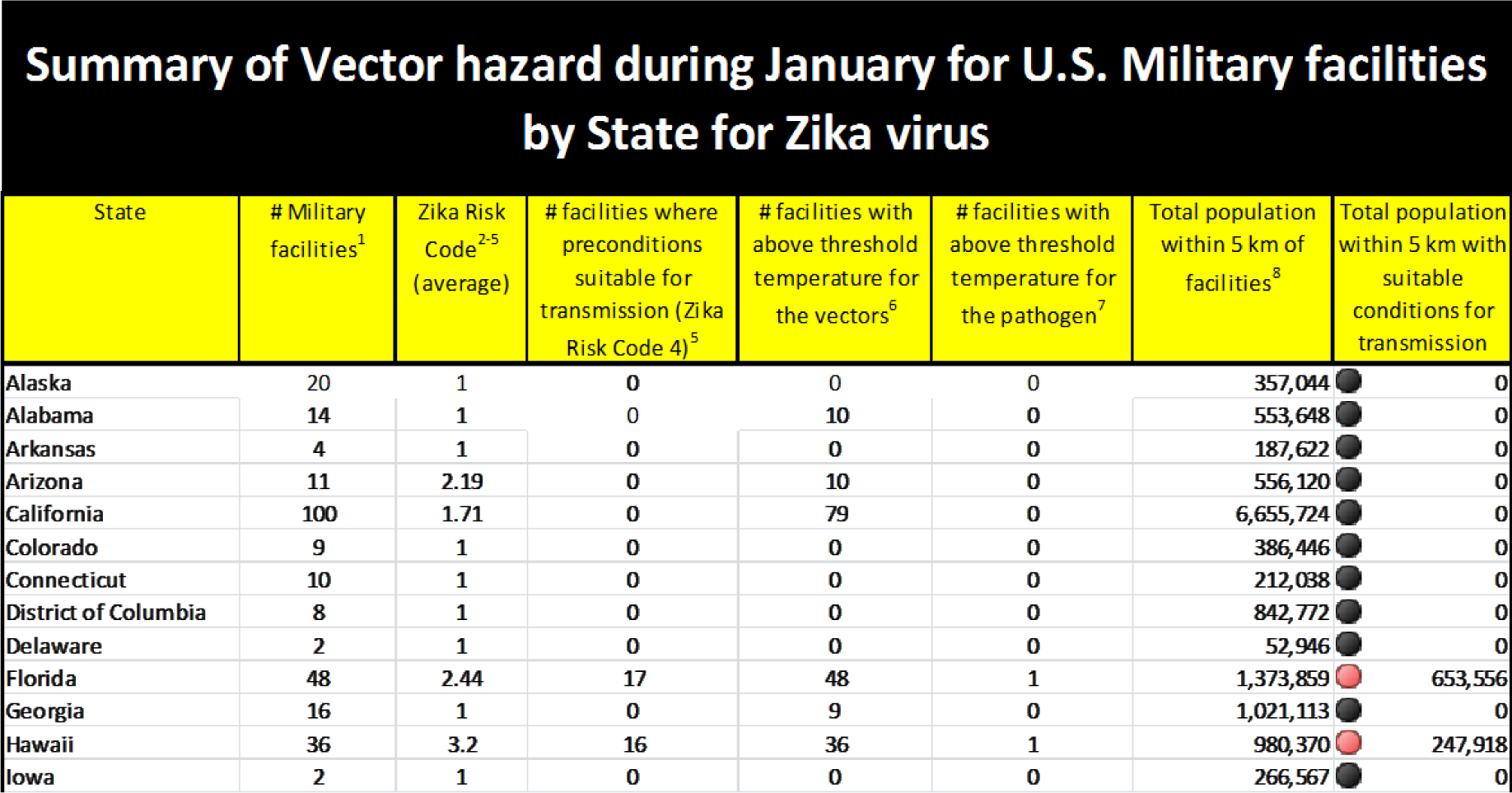
Summary risk data for each U.S. State to assist with public health and resource allocation planning.

**Figure 7.**
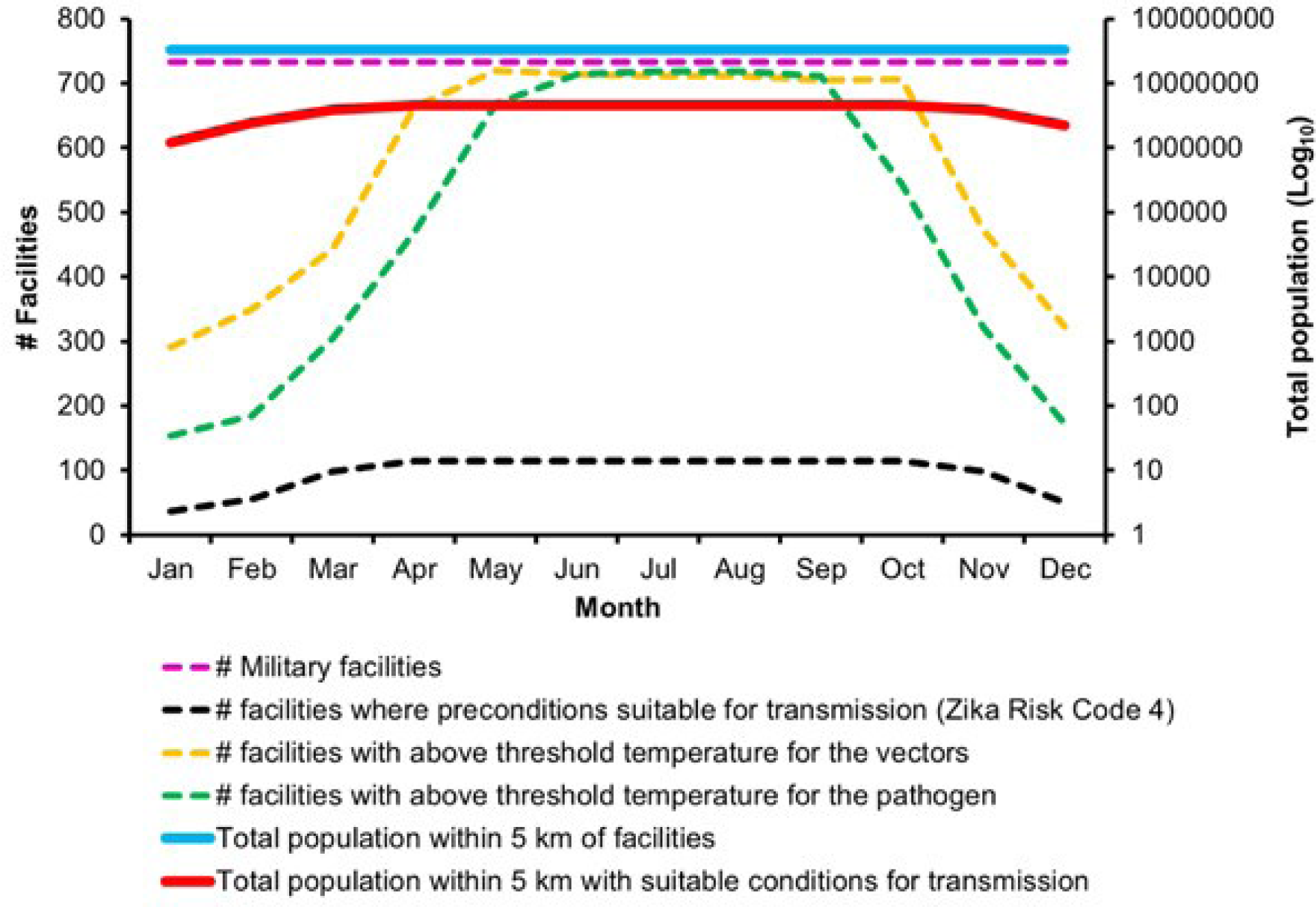
Summary risk data for 733 U.S. military facilities over 12 months.

These data may assist public health planning, and can be seen as an indicator of potential disease burden, or of people potentially benefiting from a well-informed vector surveillance and control program conducted within military facilities. Results are provided in a variety of symbologies and as textualized statements of how the factors examined may impact ZIKAV transmissions, and recommended actions for entomologists conducting routine vector surveillance. The action statement textualizes the data and is designed to assist a preparedness posture particularly around vector surveillance and control. Changes in the action statement over the year, for example as a result of rising temperature, can be used as a guide to affect changes in vector surveillance and control activities at particular facilities.

## DISCUSSION

This Excel tool is designed to provide insights into ZIKAV transmission potential at U.S. military facilities, but could be applied to other arboviruses and situations, such as cities (Monaghan et al. 2016), tire dumps or parks. The spreadsheet is flexible in that vector and virus suitability model scores, temperature limits, and the wording of action statements can be replaced depending on the context, and as new information comes to light.

For U.S. military situations, this tool could be used in conjunction with the Electronic Surveillance System for the Early Notification of Community-based Epidemics (ESSENCE) or Medical Situational Awareness in Theater (MSAT), which reports on febrile illnesses and rash in the military population. Coordination of result reporting through the Armed Forces Pest Management Board (AFPMB) and VectorMap may also be desirable. The Navy and Marine Corps Public Health Center’s guide (NMCPHC, 2016) states that “each installation’s medical personnel should conduct ongoing *Aedes* surveillance during the mosquito season appropriate to their region and take preventive and responsive action to reduce disease risk to active duty, government employees, and family member populations”. In addition, the DoD instruction OPNAVINST 6250.4C “requires all Navy and Marine Corps installations to have an Emergency Vector Control Plan (EVCP) for disease vector surveillance and control during disease outbreaks”. The spreadsheet described in this study should complement “installation pest management plans, including the EVCP, as a way to assess the risk of vector borne diseases, and implement strategies to reduce the risk to personnel assigned to installations” (NMCPHC, 2016).

Knowing when conditions are suitable for vectors is crucial for monitoring the success or failure of any control program. Appendix C of NMCPHC (2016) consists of a chart to determine the risk of infection on an installation and when to apply vector control measures. This four level vector threat response plan relies on information about vector abundance and reports of disease transmission. We see the Excel spreadsheet risk tool as a valuable adjunct to the NMCPHC plan, as it would assist with defining the length of the mosquito season, and the judicious deployment and timing of entomological resources. Each military facility is unique, and varies in size, function, human density, and suitable mosquito habitat, so not all of the 733 facilities addressed in this study will be at risk of mosquito-borne disease and suitable candidates for mosquito surveillance. However, all locations, at worst, should be useful as a point of reference for other nearby locations where mosquito surveillance is conducted.

The Zika Risk Code developed here (Figure 1) derived some inspiration from Figure 3 of Fischer et al. (2013), who combined models of vector habitat suitability with temperature categories for CHIKV replication to produce a matrix of climate related risk classes.

It is important to note that each data source used in this analysis has the potential for errors which should be considered when determining risk. For example, habitat suitability models for each vector may not be accurate for all areas, and only predict average yearly suitability. Temperature data refers to the maximum day-time air temperature near the surface (averaged over various spatial resolutions) from daily data for a recent date range, which NASA acknowledges has limitations. Vectors can also seek microclimates (e.g. indoors, subterranean habitats) that may be warmer or cooler than the outside temperature that is estimated by remote sensing data. Temperatures within the suitable range may not effect organisms uniformly. According to Westbrook et al. (2010) adult females reared from immature stages at 18°C, were six times more likely to be infected with CHIKV than females reared at 32°C. Westbrook et al. (2010) noted that climate factors, such as temperature, experienced at the larval stage, which would not be detected by adult trapping programs, can influence the competence of adult females to vector arboviruses.

We also do not account for temperature fluctuations; according to Lambrechts et al. (2011), mosquitoes lived longer and were more likely to become infected with DENV under moderate temperature fluctuations, than under large temperature fluctuations. Thangamani et al. (2016) and Ferreira-de-Brito et al. (2016) found that ZIKAV can be vertically transmitted in *Ae. aegypti* but not *Ae. albopictus*. This capability suggests mechanisms for the virus to survive in eggs that can survive for months in a dried dormant state during adverse conditions, e.g. a harsh winter that would normally kill adults.

The risk levels calculated in the spreadsheets deliberately uses simplified assumptions about temperature and does not consider precipitation, interspecific competition, anthropogenic factors such as imported cases, built-up areas, vegetation indices, and economic indices that can modify risk in complex and less understood ways. It is recommended that a level of caution be taken when interpreting the data provided by this system. It is wise to monitor activity in surrounding facilities and any reputable information from other sources before acting on any recommendations given here. It is further recommended that the near-real time and forecast analysis should be viewed in conjunction with the monthly average Excel vector hazard files which uses average monthly maximum temperature, to gain further longer term insights into where thermal conditions will support vector activity.

## ACKNOWLEDGMENTS

We would like to thank Jean-Paul Chrietien (Armed Forces Health Surveillance Branch) for constructive feedback during the development of the Excel Risk Tool, as well as Jim Writer and Penny Masuoka for providing references and feedback useful in the development of the ZIKAV threshold information. We would also like to thank LTC (Ret.) Jason Richardson, LTC Jeff Clark, CDR Stell and Terry Carpenter of the Armed Forces Pest Management Board for their feedback. This study was made possible with funding from the U.S. Department of Defense Global Emerging Infections Surveillance and Response System, Silver Spring, MD (http://www.health.mil/Military-Health-Topics/Health-Readiness/Armed-Forces-Health-Surveillance-Branch/Global-Emerging-Infections-Surveillance-and-Response). The published material reflects the views of the authors and should not be construed to represent those of the Department of the Army or the Department of Defense.

